# Planarian dorsoventral Netrins control a muscle midline signaling center and regulate blastema formation

**DOI:** 10.1101/2022.08.31.506052

**Authors:** Erik G. Schad, Christian P. Petersen

**Affiliations:** Department of Molecular Biosciences, Northwestern University, Evanston, IL 60208; Robert Lurie Comprehensive Cancer Center, Northwestern University, Evanston IL 60208

## Abstract

Integration of positional information across body axes is likely critical for whole-body regeneration to define the territories of missing tissue in three dimensions with fidelity. The body-wall musculature in planarians expresses patterning factors regulating the anteroposterior, dorsoventral, and mediolateral axes, but how this information coordinates is not fully understood. We identify a previously described factor specifically expressed in dorsal midline muscle as a BMP/Activin decoy receptor *bambi-2*. Analysis of scRNAseq indicates *bambi-2+* cells coexpress midline-specifying transcription factor *pitx* and longitudinal muscle-specifying factor *myoD*, and production of *bambi-2+* cells requires these factors. In laterally amputated animals regenerating an entirely new midline, *bambi-2+* cells are initially formed at the wound site, then dynamically spread, and ultimately reset to restore bilateral symmetry. We further identify a system of dorsoventral Netrin and Netrin receptor signals expressed from body-wall muscle that control midline identity and blastema morphology. Ventral and laterally expressed *netrins -1, -4,* and *-5* signal via dorsally-enriched netrin repulsion receptors *unc5-C, unc5-E,* and *dcc-2*, which together limit mediolateral spread of *bambi-2+* dorsal midline muscle and influence the architecture of the muscle system. Our results suggest a model in which ventral determinants dictate mediolateral information important for blastema morphology.

## Introduction

Whole body regeneration involves tissue restoration from a variety of possible truncations to axis content after amputation. Species of planarians, acoels, and cnidarians exemplify these abilities, as they are able to regenerate essentially entirely new animals after even severe injuries such as decapitation (Reddien, 2018; Vogg et al., 2019; Srivastava, 2022). The nature of robust positional information is therefore important to understand how extreme regenerative ability is possible. The information systems controlling regenerative ability across each body axis have often been considered separately, but surgical injuries also frequently damage more than one orthogonal body axis. How overall patterning across axes is accomplished is incompletely understood and may be important for understanding how regenerative organisms correctly match blastema specification to the identity and content of pre-existing tissue

Flatworm planarians can regenerate from nearly any type of surgical injury to restore axis content along their anteroposterior (AP), dorsoventral (DV), and mediolateral (ML) axes. A system of posterior Wnts and anterior Wnt inhibitors are expressed in gradients along the AP axis, and inhibition of canonical Wnt signaling results in ectopic head regeneration while over-activation due to inhibition of anti-Wnt factors results in ectopic tail regeneration (Gurley et al., 2008; Iglesias et al., 2008; Petersen and Reddien, 2008; Petersen and Reddien, 2009; Gurley et al., 2010; Petersen and Reddien, 2011; Owen et al., 2015; Reuter et al., 2015; Stuckemann et al., 2017). In addition, two Wnt/FGFRL systems control head and trunk regionalization in conjunction with Disheveled and Src signals (Cebria et al., 2002; Almuedo-Castillo et al., 2011; Lander and Petersen, 2016; Scimone et al., 2016; Bonar et al., 2022). By contrast, the dorsoventral (DV) axis is controlled through dorsal *bmp4* signals and ventral *admp*, which signal in a negative feedback loop (Molina et al., 2007; Orii and Watanabe, 2007; Reddien et al., 2007; Gavino and Reddien, 2011). Finally, medio-lateral axis patterning is regulated by medially expressed *slit* which opposes laterally expressed *wnt5* (Cebria et al., 2007; Gurley et al., 2010).

The major signals specifying AP, DV, and ML pattern in planarians are expressed from subepidermally located muscle body-wall cells that surround the inner parenchymal region containing the neoblast planarian stem cells (Witchley et al., 2013). Neoblasts are pluripotent and generate all planarian adult differentiated cells, thus underlying the animal’s ability to replace diverse missing tissues and undergo perpetual tissue homeostasis in the absence of injry (Wagner et al., 2011; Reddien, 2018; Zeng et al., 2018). Planarian muscle cells are mononucleated and project contractile fibers across several orientations (longitudinal, diagonal, circular, and dorsoventral orientations) to form a meshwork plexus (Witchley et al., 2013; Cebria, 2016). The majority of patterning factors controlling regionalization are expressed from multiple muscle subtypes (Scimone et al., 2017). However, analysis of cell fate transcription factors required for producing individual muscle subtypes found that some patterning genes have expression restricted to muscle oriented in particular directions. For example, injury triggers expression of the Wnt inhibitor *notum* and the Activin pathway modulator *follistatin*, important for wound polarization and outgrowth (Petersen and Reddien, 2011; Gavino et al., 2013; Roberts-Galbraith and Newmark, 2013; Tewari et al., 2018), and inhibition of the longitudinal muscle fate transcription factor *myoD* prevents their expression and leads to failed regeneration (Scimone et al., 2017). Likewise, circular muscle requires *nkx1.1* for its production, selectively expresses *activin-2* involved in polarity determination (Cloutier et al., 2021), and inhibition of *nkx1.1* leads to splitting the anterior pole and formation of split-axis animals (Scimone et al., 2017). In addition, medial dorsoventral muscle controlled by *foxF1* and *gata4/5/6-2* factors is a source of *slit* (Scimone et al., 2018). However, little is known about how the muscle system becomes organized and how axis patterning systems communicate with each other to result in a restoration of form following injury.

Prior studies have suggested the existence of communication across the body axes. The master regulator of posterior identity *wnt1* is required for posterior regeneration and is expressed selectively in the posterior along the dorsal midline (Petersen and Reddien, 2009; Gurley et al., 2010). *wnt1* is co-expressed in a posterior subpopulation of dorsal midline (DM) muscle cells which express the previously unannotated gene *dd23400* and extend across the majority of the AP axis (Fincher et al., 2018; Schad and Petersen, 2020). Overactivation of *wnt1* in *dd23400+* DM muscle cells through inhibition of *mob4* or *striatin* components of the STRIPAK complex results in the expansion of posterior tissue in homeostasis and regeneration, suggesting that regulation of DM muscle cells is important for body patterning (Schad and Petersen, 2020). Additionally, the correct placement of the *notum+* anterior pole at the mediolateral and dorsoventral midpoint is dependent on mediolateral components *wnt5* and *slit*, as well as the dorsoventral regulator *bmp4* (Oderberg et al., 2017). These observations suggest that patterning information may communicate across orthogonal axes and also the importance of the dorsal midline in regeneration. However, the roles for dorsal midline muscle cells in regeneration are not fully understood.

In order to understand how midline cells contribute to regeneration, we analyzed single-cell RNAseq data to identify factors expressed in midline muscle. This approach identified a set of transcription factors required for DM muscle identity. We found that factors required for DM formation were also required for lateral regeneration, suggesting the importance of these cells in facilitating outgrowth during formation of a new mediolateral axis through regeneration. Furthermore, by tracking expression of midline markers, we find that the re-establishment of dorsal midline during lateral regeneration proceeds from the wound edge rather than the mediolateral center of the fragment. Using candidate approaches, analysis of scRNAseq data, and functional analysis, we furthermore implicate Netrin related signals as regulators restricting the domain of DM cells. Netrins are a class of secreted proteins known for their role in axonal guidance (Hedgecock et al., 1990; Ishii et al., 1992; Kennedy et al., 1994; Serafini et al., 1994), and play a role in axon growth and regeneration in planarians (Cebria and Newmark, 2005). Other known roles for Netrins include cell survival (Mehlen et al., 1998; Llambi et al., 2001), organ morphogenesis (Dalvin et al., 2003; De Breuck et al., 2003; Srinivasan et al., 2003; Yebra et al., 2003; Liu et al., 2004), and myotube formation (Kang et al., 2004). In order to transduce their signals, Netrins bind a variety of targets including UNC-5 (Hedgecock et al., 1990) and DCC (Chan et al., 1996; Keino-Masu et al., 1996; Kolodziej et al., 1996; Fazeli et al., 1997). There are five genes encoding Netrins in planarians, all of which have prominent expression in muscle cells and are regionally expressed across the ML and DV axes (Cebria and Newmark, 2005; Scimone et al., 2017; Fincher et al., 2018). In planarians, Netrin receptors (five UNC-5s, two DCCs) are expressed widely, including in muscle cells (Fincher et al., 2018). Because Netrins and their receptors are expressed in muscle cells regionally across the ML or DV axes like other genes controlling patterning in planarians, we hypothesized that Netrins organize key features across these body axes, in particular, DM muscle cells.

## Results

### *dd23400/bambi-2+* is expressed in dorsal midline muscle

To identify factors important for DM muscle, we sought genes expressed in this cell population. Prior single-cell RNA seq analysis identified a gene *dd23400* expressed exclusively on the dorsal midline from muscle cells. This gene was originally identified without discernable sequence similarity to other proteins (Fincher et al., 2018). However, we noticed that PSI-blast of *dd23400* recovered several homologs of a family of Bmp/Activin membrane-bound inhibitors (BAMBI) (Figure S1) whose members possess extracellular ActR domains, lack intracellular activation domains, and may therefore function as decoy receptors (Onichtchouk et al., 1999). SMART domain analysis weakly identified an ActR domain in *dd23400* and also a transmembrane domain. Using maximum-likelihood phylogenies, *dd23400* placed with strong support within a clade of bilaterian BAMBI homologs separate from BMPR1 receptors, and so we named the gene *bambi-2*.

To examine the requirements for DM muscle cell production, we re-examined a prior whole-animal single-cell RNAseq atlas generated by DropSeq to search for factors expressed in *bambi-2+* muscle cells (Fincher et al., 2018). Because *bambi-2* has relatively weak expression, we searched for cells co-expressing *bambi-2* and the midline factor *slit,* and compared these to bulk muscle cells using the scde package for differential gene expression profiling from single-cell RNAseq data (Kharchenko et al., 2014). The resulting analysis identified several genes putatively enriched for expression in *bambi2+* midline muscle cells, including *pitx, bmp, msx, myoD, fgfr4, delta-like, TM211, nephrin, unc5-C,* and *fgfr4* (Figure 1A, Table S1). A prior analysis using Smart-Seq to find transcriptome information for *wnt1+* cells identified enriched expression (log fold-change > 2) in these cells for *pitx, wif, nephrin, msx, fgfr4, bmp, myoD, unc5-C, TM211, delta-like,* and *bambi-2* (Li et al., 2019). As a further test of midline enrichment, we performed colorimetric whole-mount in situ hybridizations on several candidates to examine whether which had midline expression. We found midline-related expression for 11 out of 15 factors tested, with the remaining 4 lacking detection in the assay (Figure 1A, Figure S1). The transcription factor *pitx* was identified previously as important for expression of *wnt1* (Currie and Pearson, 2013; März et al., 2013), a master regulator of posterior formation expressed from the posterior dorsal midline (Adell et al., 2009; Petersen and Reddien, 2009; Gurley et al., 2010) and also for new expression of the midline determinant *slit* in regeneration blastemas (Currie and Pearson, 2013; März et al., 2013). Posterior *wnt1* was subsequently shown to be co-expressed in a posterior subset of *23400/bambi-2+* dorsal midline muscle cells (Schad and Petersen, 2020). We used FISH to verify that *pitx* has expression along the dorsal midline (Figure 1B). We further reasoned that other factors required for *wnt1* expression might also have expression on the dorsal midline. The transcription factor *islet* is required for expression of *wnt1* (Hayashi et al., 2011) and although our analysis of a DropSeq atlas did not identify enrichment for this gene in midline muscle, a prior analysis using SmartSeq found enrichment for *islet* expression in *wnt1+* posterior midline cells (Li et al., 2019). Using FISH, we found *islet* is expressed more broadly than *pitx,* but including dorsal midline cells in the anterior and posterior of the animal (Figure 1B).

**Figure 1.**
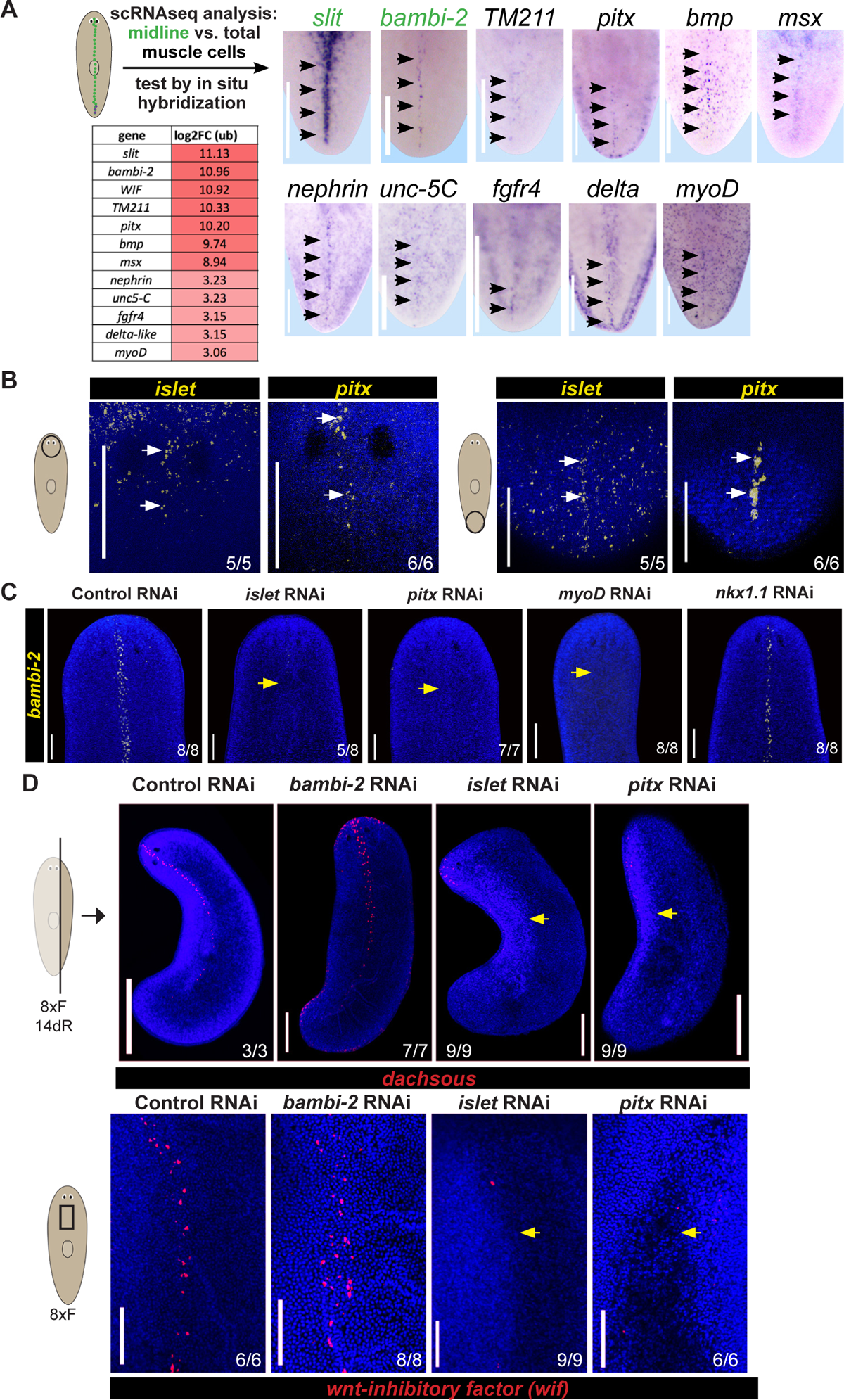
*pitx, islet,* and *myoD* are required for expression of *bambi-2*. **(A)** Analysis of prior scRNAseq data (Fincher et al 2018) comparing transcriptomes of dorsal muscle midline cells co-expressing *bambi-2(dd23400)* and *slit* versus *collagen+* muscle lacking *bambi-2* and *slit* expression. Table shows log2-fold change (upper bound, ub) from scde analysis. Right panels show WISH of candidates with expression on the posterior midline. Images represent 3-4 animals per condition. **(B)** Animals were fixed and FISH was performed to detect *islet* or *pitx* expression on the dorsal midline in the anterior (left) or posterior (right). Arrows indicate expression on the dorsal midline. **(C)** Homeostatic RNAi (8 feedings dsRNA) to inhibit transcription factors specific for midline cells (*islet, pitx*), L-muscle (*myoD*), and C-muscle (*nkx1-1*) followed by FISH to detect *bambi-2*. **(D)** FISH to detect expression of *dachsous* and *wif* midline markers after *pitx*, *islet* or *bambi-2* RNAi. Bars, 100 (B) or 200 microns (A,C).

We observed that in addition to expression on the dorsal midline in the posterior, *islet* and *pitx* are also expressed on the DM in the anterior (Figure 1B), so we reasoned that they could be required for all *bambi-2+* cells. We performed RNAi of *islet* and *pitx,* and found a nearly complete loss of *bambi-2* expression (Figure 1C). *bambi-2+* DM muscle cells may additionally be specified by transcription factors required for other muscle cell subsets. The *myoD* transcription factor is required for longitudinal muscle throughout the body (Scimone et al., 2017), was recovered by scRNAseq analysis as enriched in *bambi-2+* midline cells, and its expression domain includes a midline-enriched region expression pattern (Figure 1A). Therefore, we tested whether *myoD* inhibition would impact *bambi-2* dorsal muscle expression. RNAi of *myoD*, but not the *nkx1.1* transcription factor required for circular muscle, caused a loss of *bambi-2* expression, suggesting that *bambi-2+* DM muscle cells are likely a subset of longitudinal muscle (Figure 1B). Because *pitx* and *islet* are known to be required for expression of *slit* (Currie and Pearson, 2013; März et al., 2013), these results argue these factors likely control midline muscle cell identity in general. We tested other markers of midline to confirm these findings. *dachsous* has midline expression (Vu et al., 2019) and was enriched for expression in midline muscle by the scde analysis (Table S1, log2 upper-bound enrichment of 3.10). *pitx* or *islet* RNAi resulted in a loss of *dachsous* expression (Figure 1D). Furthermore, a homolog of *Wnt inhibitory factor (wif)* (Hsieh et al., 1999), itself a homolog of Drosophila *shifted* (Glise et al., 2005), was recovered as strongly enriched in *bambi-2+* muscle cells from the single-cell RNAseq analysis (Figure 1A, log2 upper-bound enrichment of 10.92). FISH analysis revealed *wif* to be expressed weakly along the dorsal midline, and inhibition of *pitx* or *islet* similarly reduced *wif* expression. Therefore, *pitx* and *islet* are important for expression of several markers of dorsal midline muscle. We also used the additional markers of midline muscle to examine the consequences of *bambi-2* inhibition on these cells. *bambi-2(RNAi)* animals appeared phenotypically normal and did not fail to express midline *dachsous* or *wif* (Figure 1C-D). Additionally, co-inhibition of another previously reported BAMBI homolog, *bambi,* also resulted in animals with normal appearance. Together, *bambi-2* is a marker of dorsal midline but likely not critical for its formation.

The high specificity of expression of *bambi-2* for the dorsal midline enabled the opportunity to examine the process of midline re-establishment during lateral regeneration. The expression of *bmp4* has been used to examine midline formation during lateral regeneration previously. In *Schmidtea mediterranea*, *bmp4* is expressed both on the dorsal midline as well as laterally on the dorsal side in a graded manner (Reddien et al., 2007), and in *Dugesia japonica, bmp* expression is prominent on the midline and also detectable in a graded fasion from the dorsal midline (Orii, 1998). In *Dugesia japonica* animals undergoing lateral regeneration, *bmp4* expression punctate and broad at 1 day post amputation, which could either reflect pre-existing or new expression of *bmp4* (Orii, 1998). By 3 days of regeneration, expression primarily appeared as a line running anteroposteriorly and located asymmetrically on the fragment and closer to side forming a regeneration blastema. As regeneration proceeded, the most abundant *bmp4* signal still appeared to be asymmetric in the animal but located closer to the left-right midpoint of the fragment. Similar asymmetry in *bmp4* expression early in regeneration has been reported in *Schmidtea mediterranea* with some differences. In lateral regeneration in that organism, new expression of *bmp4* was detected at 12 hours in a punctate pattern close to the wound site (Reddien et al., 2007). By 48 hours, expression was stronger on the blastema side of the fragment but without yet a discernable midline component, and midline expression could be detected by 14 days. Other tests of *bmp4* expression in lateral regeneration have detected greater expression early in regeneration (2-4 days) on the blastema side of fragments (Gavino and Reddien, 2011). Together, these prior observations suggest midline formation is unlikely to proceed through a process that places new midline at the geometric midpoint. However, because *bmp* expression is graded and includes non-medial dorsal cells, the exact nature of midline regeneration is still not fully understood. We tested the expression of *bambi-2* during a timecourse of lateral regeneration in asymmetrically laterally amputated fragments that unambiguously initially lacked midline expression. *bambi-2* expression appeared by 24 hours at the wound edge at a location furthest from the existing lateral-most domain. *Bambi-2* expression retained dorsal specificity throughout the timecourse, so this position would effectively mark the dorsal-most position in the fragment remaining following the amputation (Figure 2). Over time, this expression domain expanded while retaining dorsal specificity (100% of animals only had expression on one DV side of the animal, N at least 8 animals per timepoint), reached a point of largest mediolateral spread by 72 hours, followed by restriction to a location between the pre-existing tissue and growing blastema. Consistent with prior studies, midline expression had not become fully restored to the middle of the fragment by 7 days. These results support that dorsal midline regeneration proceeds initially in close association with the wound site, followed by subsequent re-establishment.

**Figure 2.**
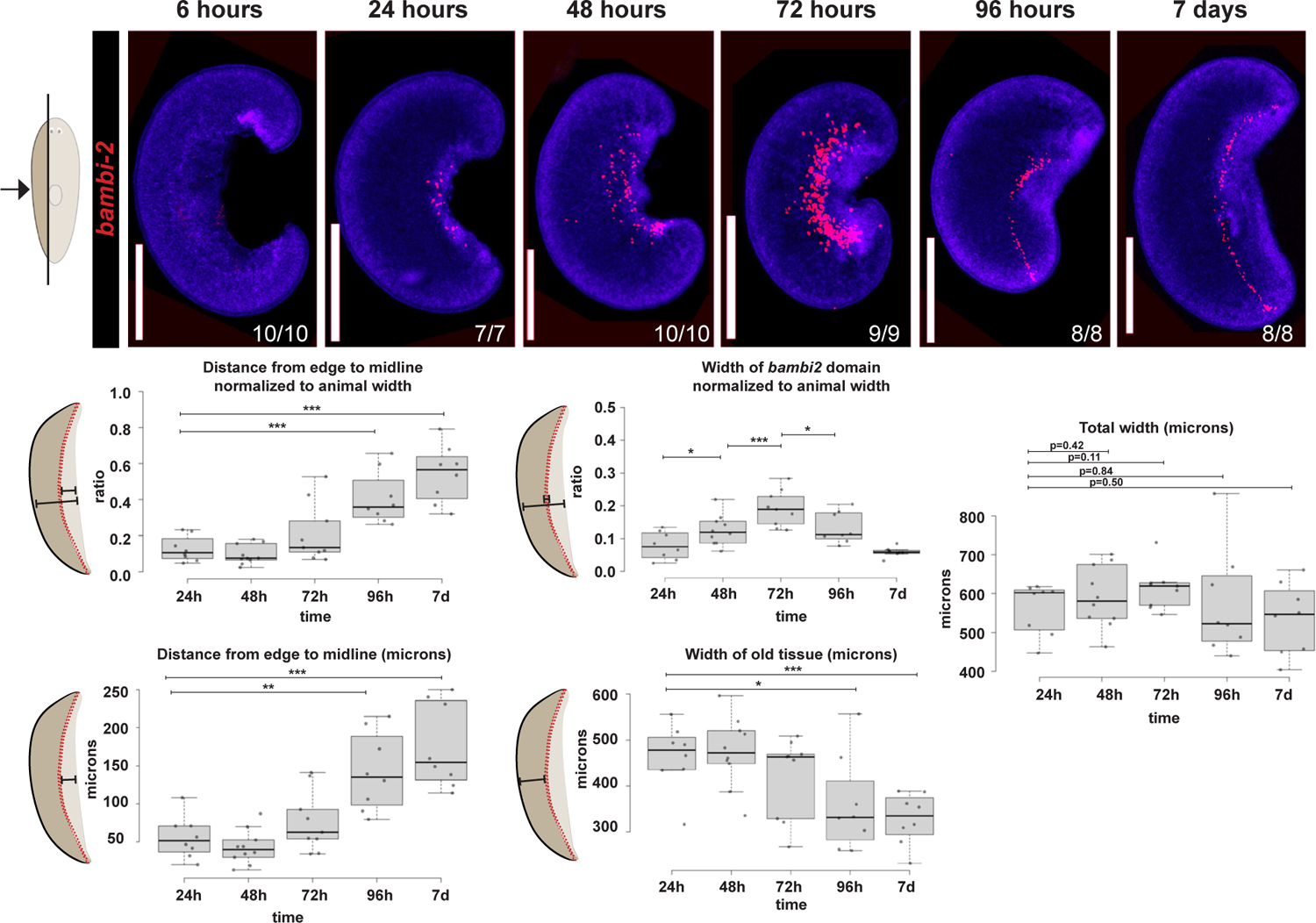
Lateral regeneration progressively restores *bambi-2* midline expression through injury-induced expression and growth. FISH of *bambi-2* expression during a timeseries of the regeneration of asymmetric lateral fragments that must regenerate the dorsal midline. Bottom, quantifications of the relative and absolute locations of the *bambi-2+* cells with respect to the wound edge (left), the relative size of the *bambi-2+* domain and size of pre-existing tissue domain lateral to the *bambi-2* domain (center), and absolute width of the regenerating fragments. *bambi-2* expression first arises at the lateral wound edge by 24 hours, followed by a progressive relative movement toward the left/right center of the fragment, rather than a mechanism that directly places these cells centrally. n=12-15 animals. * p<0.05, **p<0.01, ***p<0.001 by two-tailed unpaired t-tests.

### Netrins from ventral/medial muscle domains coordinate to instruct blastema outgrowth shape

We hypothesized from these results that dorsoventral signals could be important for establishing the *bambi-2+* expression domain. We noticed that one of the genes enriched in midline muscle expression from the scRNAseq analysis (*unc5-*C) belonged to the Netrin signaling pathway, suggesting a possible role for this pathway in DM muscle function. Netrins are secreted proteins that provide guidance cues for migratory cells (Hedgecock et al., 1990; Ishii et al., 1992) and can signal to receiving cells across long ranges (Colavita and Culotti, 1998; Hong et al., 1999; Keleman and Dickson, 2001; Finci et al., 2014; Boyer and Gupton, 2018). The planarian genome contains five genes encoding Netrin proteins, each expressed in muscle cells (Figure 3A-C, Figure S2) (Fincher et al., 2018), while *netrin-1* and *netrin-2* are also expressed in neurons (Cebria and Newmark, 2005; Scimone et al., 2017). Planarian Netrins are expressed in a series of ML and DV domains (Figure 3A). *netrin-1* and *netrin-3* are coexpressed in *collagen+* muscle cells on the lateral edge of the animal, while *netrin-2* and *netrin-5* are expressed in medial muscle cells on the ventral side (Figure 3A). *netrin-4* is expressed across the ventral side of the animal (Figure 3A). Sections of animals co-stained with each Netrin and the muscle marker *collagen* showed that Netrins are expressed in several ventral/medial domains within muscle. Because Netrins function in cell guidance and have graded expression in muscle cells on the ML axis, we hypothesized that they play a role in organizing the DM muscle domain. The planarian genome encodes several Netrin receptors: five putative UNC-5 receptors, and two DCC receptors (Scimone et al., 2020). Each of these receptors is expressed broadly, but also has graded expression in muscle cells (Fincher et al., 2018) (Figure 3B-C, Figure S2). *unc5-A,-B,-D* and *dcc-1/net-R* are expressed on the lateral edge of the animal, while *unc5-C,-E* and *dcc-2* have prominent expression in dorsal medial *collagen+* muscle cells (Figure 3B-C). In summary, Netrins are expressed in either lateral (*netrin-1,-3),* ventral-medial (*netrin 2,-5*), or broad ventral (*netrin-4*) domains, while putative receptors are expressed in lateral (*unc5-A,-B,-D* and *dcc-1/net-R*) or dorso-medial (*unc5-C,-E* and *dcc-2*) domains (Figure 3D).

**Figure 3.**
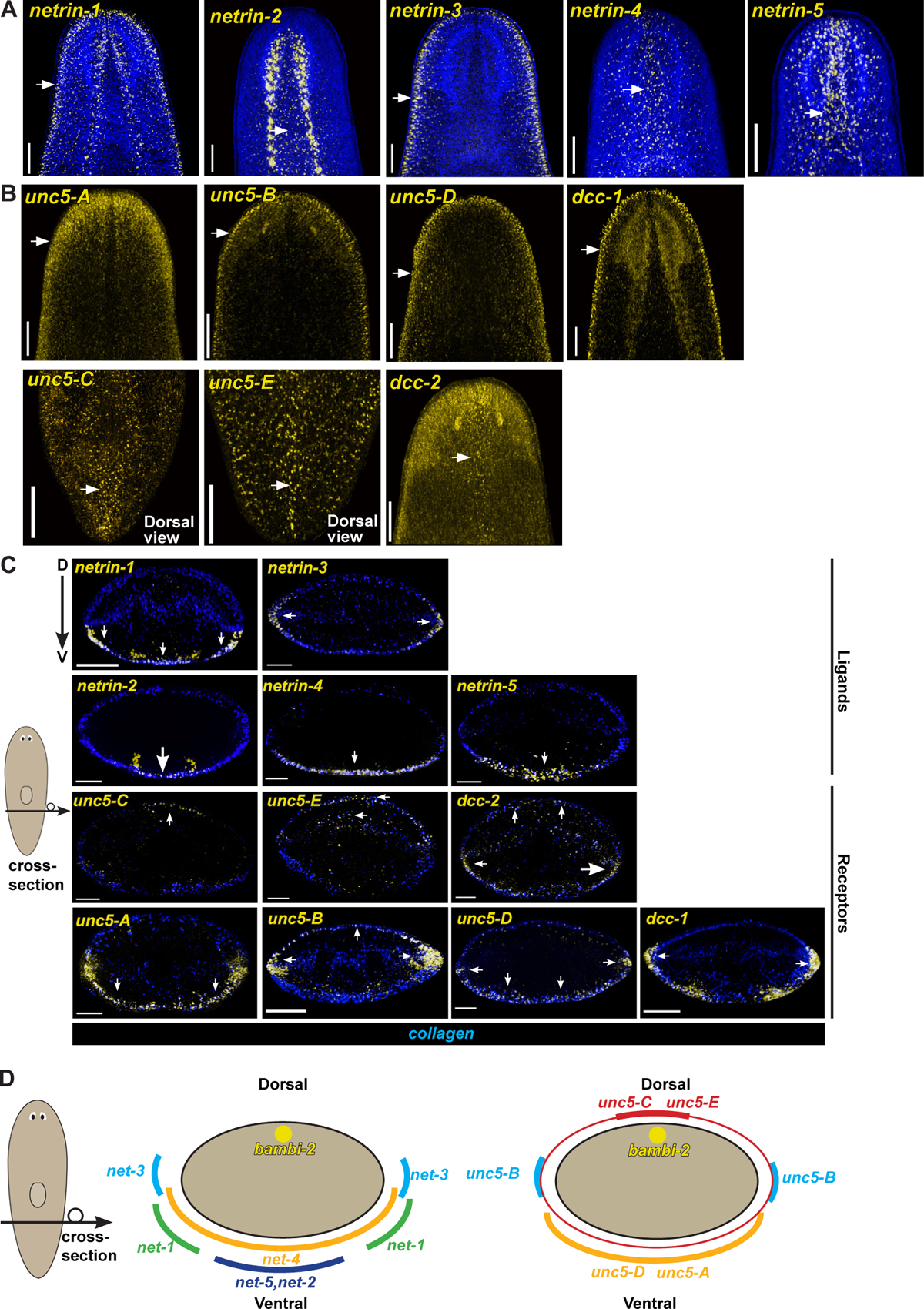
Netrins and Netrin receptors are expressed in dorsoventral domains in muscle cells. **(A)** FISH images showing *netrins* are expressed in along either ventromedial (*netrin-2, netrin-4, netrin-5)* or lateral (*netrin-1, netrin-3)* domains. Each image shows a ventral view. N=5-6 animals. **(B)** FISH images showing Netrin receptors are expressed in along either dorsomedial (*unc5-C, unc5-E, dcc-2)* or lateral (*unc5-A, unc5-B, unc5-D, dcc-1)* domains (4-7 animals probed per condition). (C) Images of cross-sections, dorsal side top. FISH images showing *netrins* are expressed in along either ventromedial (*netrin-2, netrin-4, netrin-5)* or lateral (*netrin-1, netrin-3)* domains (4-6 animals probed per condition) FISH images showing Netrin receptors are expressed in along either dorsomedial (*unc5-C, unc5-E, dcc-2)* or lateral (*unc5-A, unc5-B, unc5-D, dcc-1)* domains. Bars, 100 (A,B) or 50 microns (C). Arrows indicate expression of each tested gene coincident with *collagen+* muscle. (D) Cartoon of planarian cross-section displaying the approximate expression domains for each Netrin and Netrin receptor.

To investigate the role of Netrins during regeneration we inhibited each Netrin using RNAi. We first inhibited each Netrin individually, resulting in no overt phenotypes of dysfunction in either anterior or posterior regeneration (Figure 4A). We reasoned that Netrins could function redundantly, and so we inhibited all five planarian Netrins through combined RNAi, and verified by qPCR for effective knockdown (Figure S3). Inhibition of all Netrins simultaneously resulted in regeneration with midline defects of a pointy blastema in the anterior (32/36 regenerating trunk fragments), and of indented blastemas in the posterior (7/36) (Figure 4B). In addition, at a low frequency, inhibition of the five Netrins resulted in failed anterior regeneration in amputated tail fragments (6/36) or failed posterior regeneration from amputated head fragments (5/36 animals). We reasoned that these phenotypes were either due to inhibition of all Netrins together or instead due to inhibition of subsets of Netrins. To test this hypothesis, we inhibited each possible combination of two, three, or four Netrins, and scored animals for the penetrance and expressivity of pointy blastema in anterior blastemas or indentations in posterior blastemas (Figure S4A-B). We found that inhibition of all combinations of Netrins resulted in anterior pointy blastemas to varying degrees. Inhibition conditions including RNAi of *netrin-4,-5* yielded the strongest effects on pointy blastemas, followed by those including *netrin-1* and *netrin-2*. The laterally expressed *netrin-3* factor likely did not contribute, because *netrin-1;-2;-3;-4;-5* RNAi yielded the same outcome as *netrin-1;-2;-4;-5* RNAi (Figure S4A). Therefore, *netrin-1, −2,-4,* and *-5* contribute to normal anterior regeneration morphology.

**Figure 4.**
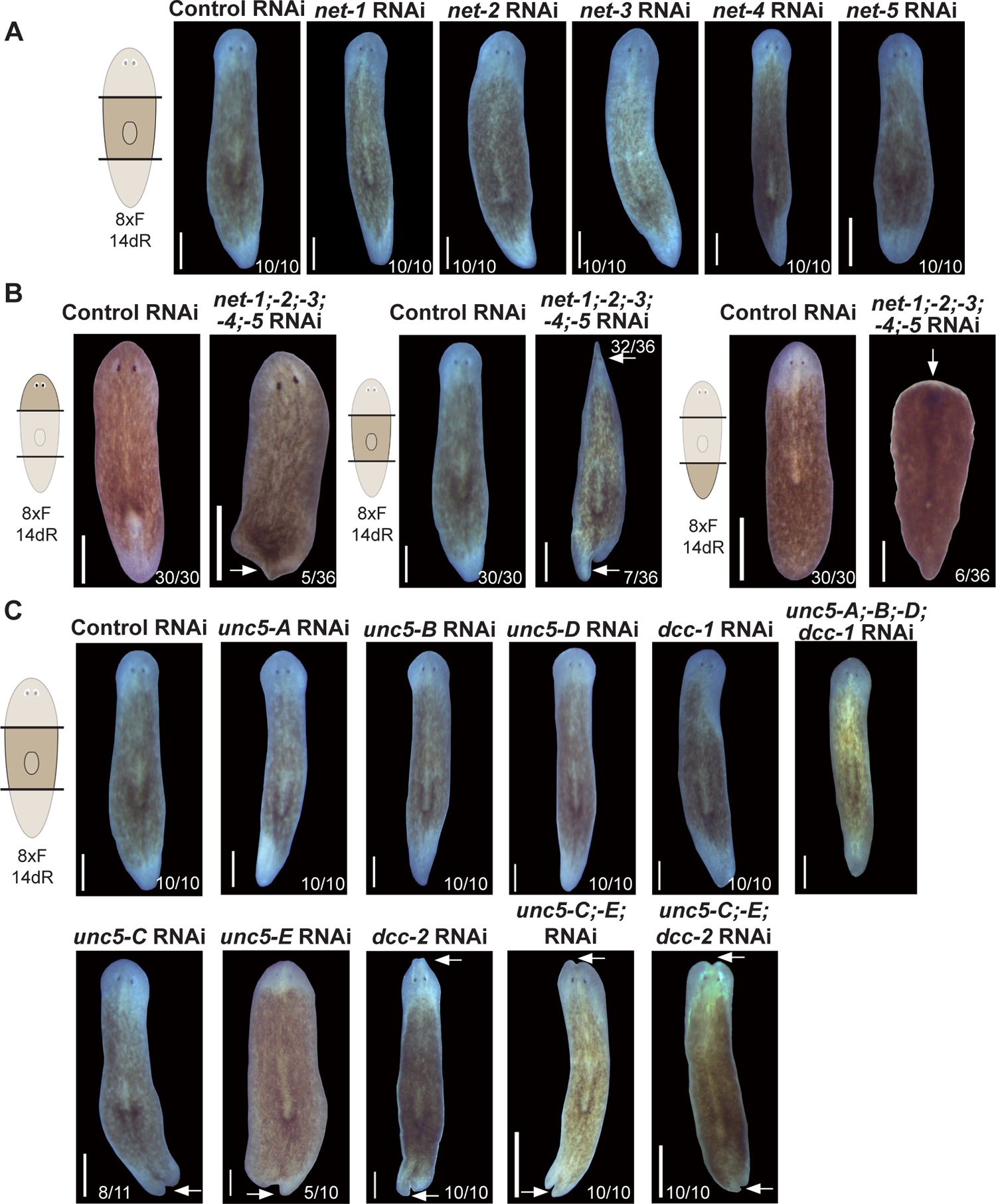
Netrins act cooperatively to control blastema morphology and regeneration. **(A)** Live images showing RNAi of individual *netrins* results in normal regeneration. (10 animals per condition). **(B)** Simultaneous inhibition of all five Netrins causes regeneration of blastemas with an indentation in the posterior and pointed morphology in the anterior, as well as weakly penetrant regeneration failure in head and tail fragments (arrows). **(C)** RNAi of *unc5-C, unc5-E* and *dcc-2* individually or in combination results in indented blastemas (arrows). Bars, 200 microns.

A similar analysis of the indented blastema phenotype from posterior-facing wounds found a somewhat distinct outcome. Whereas inhibition of all five Netrins simultaneously resulted in an indented blastema, inhibition of most combinations of Netrins resulted in normal posterior regeneration. Inhibition of *netrin-1, −4,* and *-5* increased the penetrance and expressivity of indentation over that from inhibition of all netrins, indicating a particular importance for these three genes in enabling midline regeneration in the posterior (Figure S4B). Consistent with this model, RNAi combinations excluding inhibition of any of these three Netrins resulted in normal posterior regeneration, and co-inhibitions with *netrin-2* or *netrin-3* did not strongly affect the outcome. Therefore, *netrin-1*, *netrin-4*, and *netrin-5* act together to ensure midline regeneration in blastemas generating posterior tissue.

We also examined whether the low frequency of failed regeneration after inhibition of all Netrins simultaneously could be explained by lack of injury-induced gene expression from muscle cells. However, *netrin-1;-2;-3;-4;-5(RNAi)* animals had normal injury-induced expression of *notum* and *follistatin* (Figure S5). We suggest that the failed regeneration phenotype could either reflect a distinct role for Netrins or a more extreme version of the defects causing impaired blastema morphology. Nonetheless, these results suggest that Netrin inhibition does not result in failed responsiveness of longitudinal muscle cells to injury (Scimone et al., 2017). These results also suggest that the indented morphology phenotype is not likely due to a problem with early expression of injury-induced *notum* or *wnt1*.

To further investigate the role of Netrin signaling in regeneration, we conducted RNAi experiments to test for functions of the Netrin receptors. Inhibition of the receptors expressed in lateral domains (*unc5-A,-B,-D* and *dcc-1/net-R*) appeared to result in normal regeneration, including when inhibited in combination (Figure 4C). However, inhibition of either *unc5-C, unc5-E*, or *dcc-2* caused regeneration with indentations to posterior blastemas in all cases, and indentations to anterior regeneration after *dcc-2* RNAi. Dual inhibition of *unc5-E* and *unc5-C* or triple inhibition of *unc5-E;unc5-C;dcc-2* resulted in more strongly expressive anterior and posterior blastema indentations. The indented posterior blastemas were reminiscent of the posterior blastema morphology phenotypes after RNAi of *netrins-1, −4,* and *-5*, suggesting these Netrins may signal through *unc5-E/unc5-C/dcc-2* receptors. Furthermore, while *netrin-1, netrin-4* and *netrin-5* have expression in ventral and lateral muscle domains, *unc5-E* and *unc5-C* have enrichment in a dorsal domain (Figure 3). These observations implicate a system of Netrin ligands and receptors signaling across muscle domains in to control midline blastema morphology.

### Netrins act as repulsive signals to restrict the midline domain

Based on the overt mediolateral defects in blastema formation after Netrin RNAi, we tested the expression of ML and DV patterning factors in these animals. *netrin-1;-2;-3;-4;-5(RNAi)* animals displayed an increase in *bambi-2* domain width, especially in the anterior (Figure 5A). Similarly, midline expression of *slit* was also expanded in *netrin-1;-2;-3;-4;-5(RNAi)* animals, while laterally expressed *wnt5* was unchanged (Figure 5A-C). Furthermore, *netrin-1;-2;-3;-4;-5(RNAi)* animals retained expression of dorsal *bmp4* and ventral expression of *admp*, suggesting these animals did not undergo ventralization or dorsalization (Figure 5A). Finally, *netrin-1;-2;-3;-4;-5(RNAi)* animals succeeded in expressing *wnt1+* at the posterior pole and *notum* at the anterior pole, consistent with animals regenerating with normal head and tail identity in spite of the blastema morphology defects (Figure 5A). We noted that although *notum* expression was correctly located at the anterior pole, its expression was expanded over the Netrin(RNAi) pointy blastema compared to controls. Prior studies suggest that a preexisting midline can determine the location of a newly regenerated anterior pole (Oderberg et al., 2017), so perhaps the expanded midline domain in *netrin-1;-2;-3;-4;-5(RNAi)* animals may cause formation of an expanded anterior pole. Together, these findings implicate Netrins specifically in restricting dorsal midline identity. Based on these findings, we specifically tested for any midline defects after inhibition of the Netrin receptors involved in midline formation. Simultaneous RNAi of *unc5-C, unc5-E,* and *dcc-2* resulted in a similar expansion of the *bambi-2+* cell domain (Figure 3B-C). Therefore, the phenotypes of dorsal midline expansion correlate with indented blastemas. Reception of Netrins through receptor heterodimers of Unc5 and DCC family members are thought to signal repulsion during axon guidance and cell migration (Lai Wing Sun et al., 2011). Furthermore, the *bambi-2+* dorsal midline cells are located at the point furthest away from the sources of ventral and medial *netrin-1, netrin −4* and *netrin-5* but within the dorsal domains of *unc5-E* and *unc5-C* expression. Therefore, we suggest that the *bambi-2+* cells are restricted dorsally because of repulsive signaling between ventral/lateral *netrin-1,-4,-5* to dorsally expressed Unc5 receptors *unc5-E* and *unc5-C* signaling through *dcc-2*.

**Figure 5.**
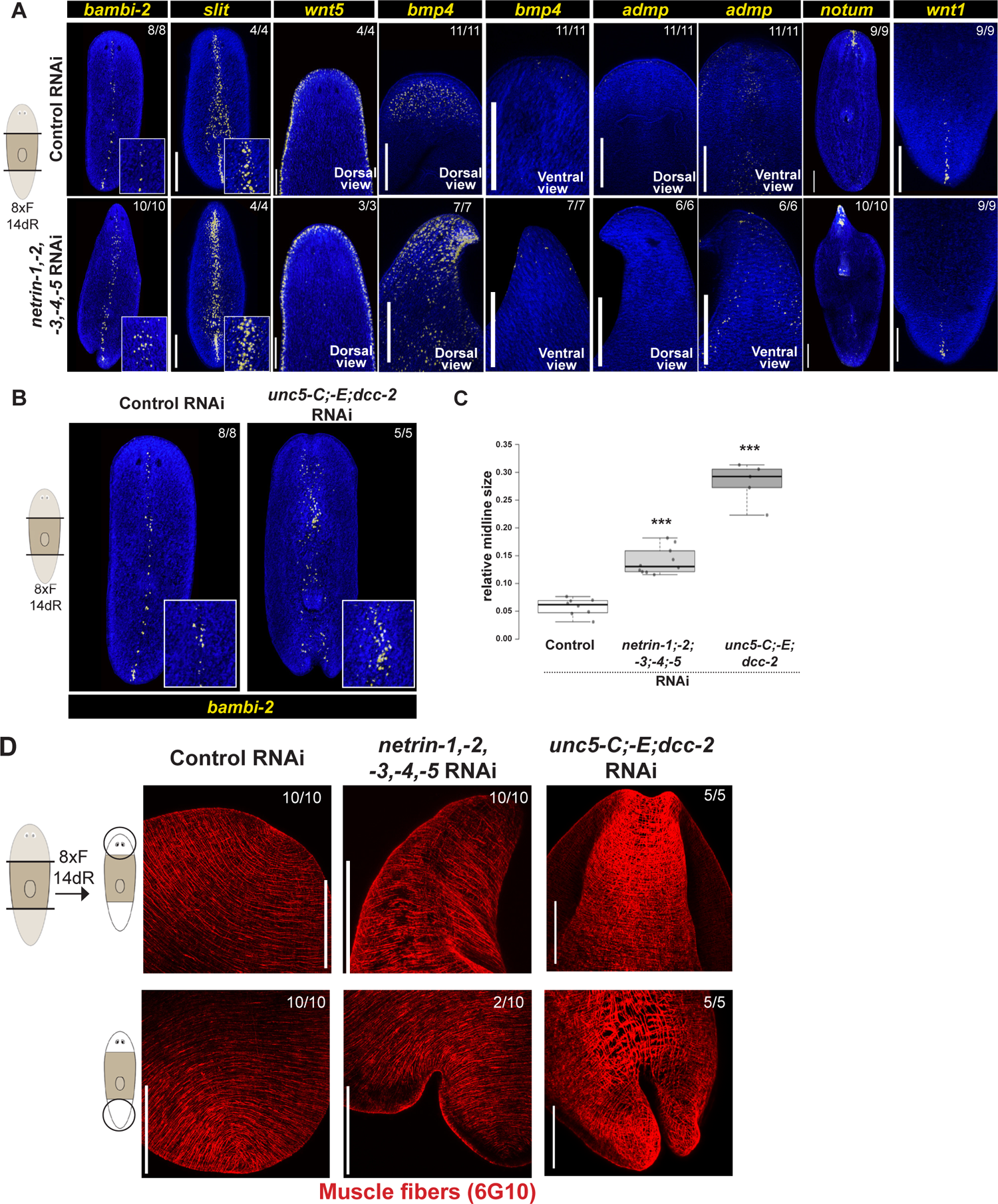
Netrins control midline muscle cell organization and proper muscle morphology. **(A)** FISH showing expression of patterning factors following inhibition of each Netrin. Midline (*bambi-2, slit*) and anterior pole cells (*notum)* are expanded, while lateral (*wnt5*), posterior (*wnt1*), and dorsoventral markers (*bmp-4, admp-1*) appear normal. *admp-* and *notum*-stained animals are from regenerating tail fragments, all other panels show regenerating trunk fragments. Insets show enlargements of *bambi-2* and *slit* expression **(B)** FISH showing expanded *bambi-2* expression domain following *unc5-C,-E,dcc-2* RNAi. Insets show enlargements of *bambi-2* expression. **(C)** Quantification of midline expansion normalized to body width in each condition. Bars show standard error of 5-10 replicates, *** p <0.001 by 2-tailed unpaired t-test. **(D)** Immunostaining with 6G10 antibody to visualize muscle fiber morphology after *netrin-1;-2;-3;-4;-5* RNAi and *unc5-C;unc5-E;dcc-2* RNAi in anterior (top) or posterior (bottom) blastemas. Muscle morphology was disrupted at locations of the indented blastema. Animals were fixed after 8 dsRNA feedings and 14 days of regeneration in all panels. Bars, 200 (A-B) or 100 microns (D).

The expression of Netrins within muscle cells led us to hypothesize that Netrins might control some aspect of organization of the muscle fiber network important for blastema formation in regeneration. To test this hypothesis, we inhibited all Netrins or Netrin receptors *unc5-E, unc5-C,* and *dcc-2*, and then used immunofluorescence to label animals with a monoclonal antibody (6G10) that labels muscle cell fibers (Ross et al., 2015) (Figure 5D). In *netrin-1,-2,-3,-4,-5(RNAi)* animals, muscle cell fibers were present in the pointed anterior blastemas and appeared normal, although they appeared to track with the overall pointed morphology. In *netrin-1,-2,-3,-4,-5(RNAi)* animals, some circular muscle fibers near the indented posterior regenerated tissue inappropriately projected anteroposteriorly after crossing the midline. *unc5-E;unc5-C;dcc-2* RNAi led to a more severe defect of indentation, and we also noticed such animals have a zone of muscle disorganization immediately anterior to the indentation along the midline, which could reflect a muscle remodeling process linked to indented tissue. We conclude that a system of ventrally expressed Netrins and dorsally expressed Netrin Receptors are likely not required for muscle fiber growth altogether, but impact the architecture of the muscle system.

## Discussion

Whole-body regeneration requires the re-establishment of patterning across all major body axes. Here, we identify processes important for generating, restricting, and using *bambi-2+* dorsal midline muscle in planarian regeneration. Inhibition of factors *pitx* and *islet* caused loss of midline expression of *bambi-2, dachsous,* and *wif* and also prevented blastema outgrowth during lateral regeneration, suggesting the importance of these cells in restoring mediolateral tissue distribution. We furthermore used *bambi-2* expression as a probe to uncover the overall process of midline regeneration from fragments lacking a midline. *bambi-2* expression initiates at the lateral wound edge. Subsequently, *bambi-2* expression expands then appears to restrict to a region between new and old tissue, and differential growth over several weeks ultimately restores bilateral symmetry (Figure 6A).

**Figure 6.**
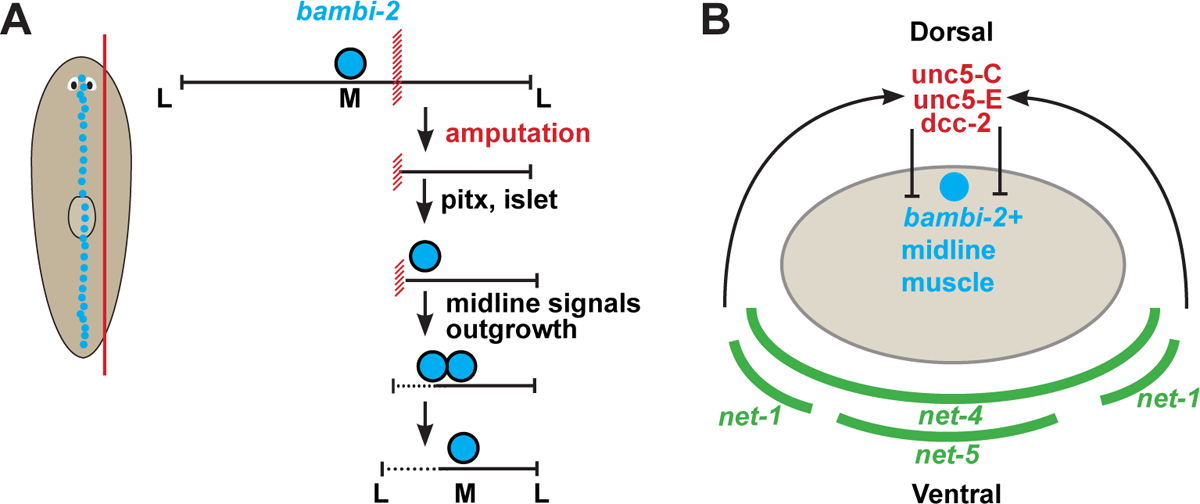
Model. **(A)** In regeneration of lateral fragments, the *bambi-2*+ dorsal midline muscle is re-established near the wound site and not the geometric center of the fragment. *pitx* and *islet* are necessary both for *bambi-2* expression and blastema outgrowth that restores bilateral symmetry (B) Three ventral netrins likely act through dorsal netrin receptors to restrict the location of the *bambi-2+* cells.

In addition, we describe the essential role of Netrin signaling in ML patterning during regeneration in planarians (Figure 6B). *netrin-1,-2,-3,-4,-5* inhibition lead to an expansion of the *bambi-2+* cell domain, but did not alter overall DV or AP identities. Netrins are known for their role in axonal growth (Hedgecock et al., 1990; Ishii et al., 1992; Hamelin et al., 1993; Colavita and Culotti, 1998), including in planarians (Cebria and Newmark, 2005), but Netrins can also guide myotube formation and other migrating cells in murine development (Dalvin et al., 2003; De Breuck et al., 2003; Srinivasan et al., 2003; Yebra et al., 2003; Kang et al., 2004; Liu et al., 2004), so it is plausible that Netrins may function to guide muscle cell growth in planarians. Three putative Netrin receptors (*unc5-C, -E, dcc-2*) are expressed in dorso-medial muscle cells, and their inhibition leads to an expansion of the *bambi-2+* cell domain. These results are consistent with a model in which ventral Netrins act as repellants by binding to these receptors (Figure 6B). When *netrin-1,-2,-3,-4,-5* are inhibited together, anterior blastemas form with a pointed morphology, with expanded *notum+* anterior pole cells. Prior work determined that the midline instructs where the anterior pole is placed in the anterior blastema (Oderberg et al., 2017), suggesting that the expanded dorsal muscle domain from Netrin RNAi conditions may cause the regeneration of an excessively wide anterior pole, thus generating pointed blastemas. Alternatively, it is possible that the pointy blastema phenotype represents a loss of other outputs for Netrin activity. For example, Netrins can function as attractants in axon guidance or migration, and this function is thought to occur via DCC/DCC receptor dimers rather than DCC/Unc5 dimers (Lai Wing Sun et al., 2011). In our survey of Netrin receptor RNAi phenotypes, no condition phenocopied the pointy blastema defect observed from Netrin RNAi. One possible explanation could be that due to the shared use of DCC in both repulsion and attraction, it is not possible to use RNAi of these factors to selectively disrupt the process of attraction. Alternatively, perhaps other receptors are the targets of Netrin signaling disrupted during the acquisition of the pointy blastema phenotype. Along these lines, we also did not find blastema failure phenotypes through inhibition of Netrin receptors. However, our phenotypic analysis identified shared functions in controlling the regeneration of midline tissue in the posterior as well as in muscle targeting, for a subset of Netrins and Netrin receptors. RNAi of *netrin-1;-4;-5* or *unc5-C;unc5-E;dcc-2* impacted the organization of the planarian musculature system, causing the newly made circular muscle in regenerating blastemas to mistarget over the midline and instead forming separated domains that grow into indented blastemas.

Whole body regeneration in planarians depends on axis information systems encoded in muscle cells. Our study finds muscle-expressed ligands signaling through muscle-expressed receptors to control regionalization of muscle identity. These observations suggest that regionalization signaling could take place directly between muscle cells. Alternatively, given that *dcc-2* and other netrin receptors are expressed more broadly, an alternative possible mechanism for pattern formation could involve the use of positional cues from muscle to influence regionalization through control of stem cell differentiation into distinct muscle domains. Netrins have dorsoventral expression domains opposing BMP in multiple organisms, including in planarians (Cebria et al., 2007; Reddien et al., 2007), hemichordates (Lowe et al., 2006), acoels (Srivastava et al., 2014), Drosophila (Harris et al., 1996; Mitchell et al., 1996), and within the vertebrate neural tube (Serafini et al., 1994; Lai Wing Sun et al., 2011). In addition, Netrins have expression in body wall muscle in planarians (Witchley et al., 2013), acoels (Raz et al., 2017), *Drosophila*, and *C. elegans* (Ishii et al., 1992). Our observations point to an ancient and conserved use of Netrins for linking dorsoventral and midline identity through muscle. The involvement of multiple Netrins and receptors for controlling dorsal midline muscle and blastema morphology suggests the possibility of a dorsoventral code of these factors that guide patterning of the blastema in order to restore bilateral symmetry during whole-body regeneration.

## Acknowledgements

We thank members of the Petersen lab for critical comments. CP acknowledges funding from NIGMS R01GM129339, R01GM130835, and pilot project funding from the NSF-Simons Center for Quantitative Biology at Northwestern University, an NSF (1764421)-Simons/SFARI (597491-RWC) MathBioSys Research Center.

## Author contributions

CP conceived of the study and acquired funding, ES designed and conducted experiments, CP and ES wrote the manuscript.

## Declaration of Interests

There are no competing interests.

## Supplemental Figures

**Figure S1.**
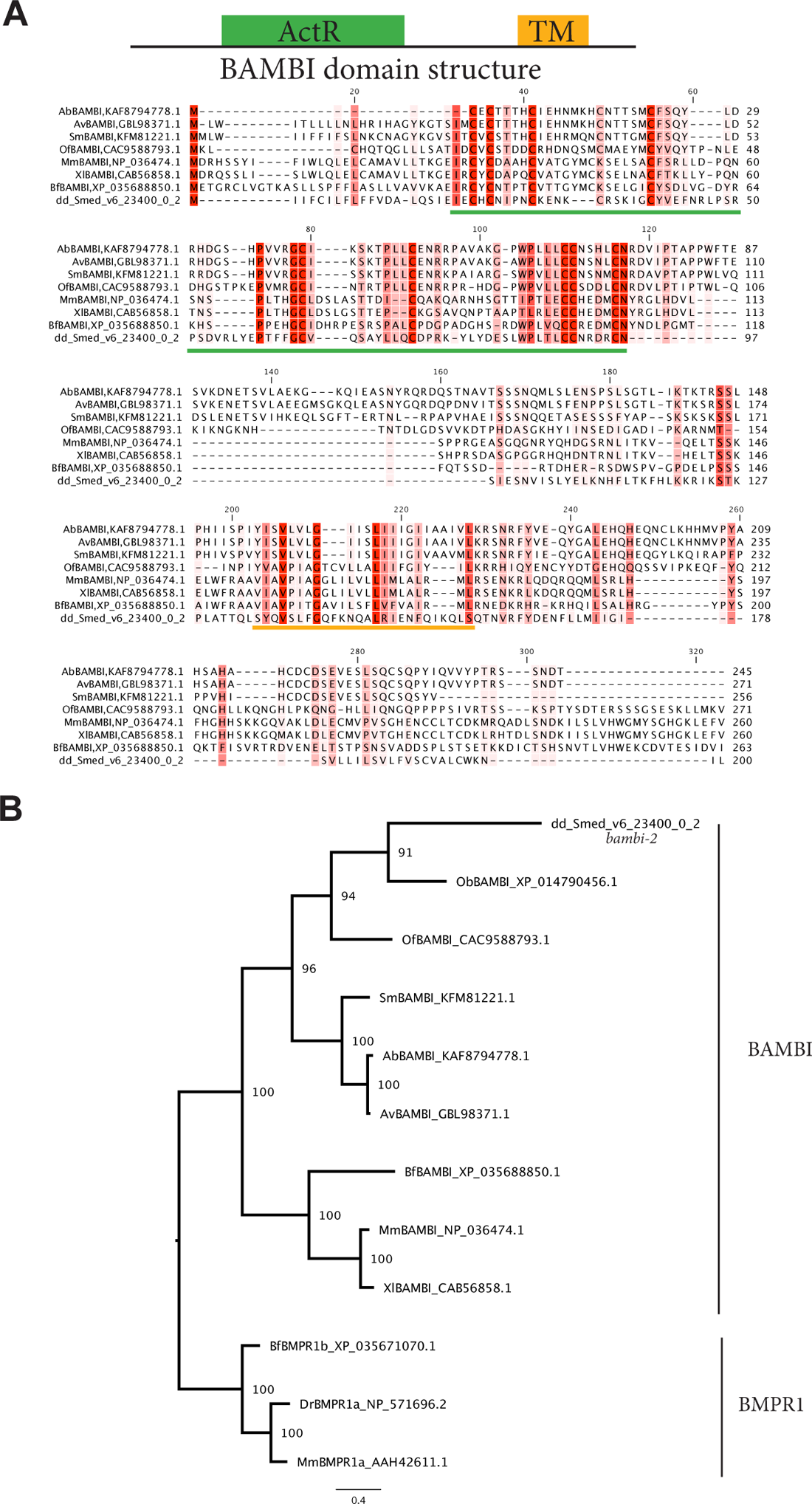
dd23400/*bambi-2* encodes a BAMBI decoy receptor. **(A)** Sequence alignment of dd23400 with BAMBI factors with conserved residues highlighted in red. Green bar indicates location of ActR domain. dd23400 has a weakly detected ActR domain (aa4-84, evalue 0.14), possesses a transmembrane domain (aa158-180, TMHMMv2.0) and like other BAMBI homologs lacks an intracellular kinase activation domain. DELTA Blast of dd23400 followed by PSI-blast (3 rounds, 500 iterations) recovers BAMBI orthologs from several species, including BAMBI from Osmia bicornis (evalue=1e-22), Megachile rotundata (evalue=3e-22) and Apis mellifera (evalue=3e-20).**(B)** Bayesian inference phylogenetic tree generated from alignment with MUSCLE and calculated with MrBayes v3.2.3 using a GTR substitution model with gamma-distributed rate variation across sites and a proportion of invariable sites. Runs of 50000 generations converged with an average standard deviation of split frequencies <0.01 and 25% of trees were discarded as burn-in. Tree displayed using FigTree v1.4.4 with branch labels indicating posterior probabilities. BAMBI sequences from Owenia fusiformis (Of, CAC9588793.1), Octopus bimaculoides (Ob, XP_014790456.1), Argiope bruennichi (Ab, KAF8794778.1), Araneus ventricosus (Av, GBL98371.1), Stegodyphus mimosarum (Sm, KFM81221.1), Xenopus laevis (Xl, CAB56858.1), Mus musculus (Mm, NP_036474.1), and Branchiostoma floridae (Bf, XP_035688850.1).Tree is rooted using BMPR1 receptors from Brachiostoma floridae (Bf), Danio rerio (Dr), and Mus musculus (Mm).

**Figure S2.**
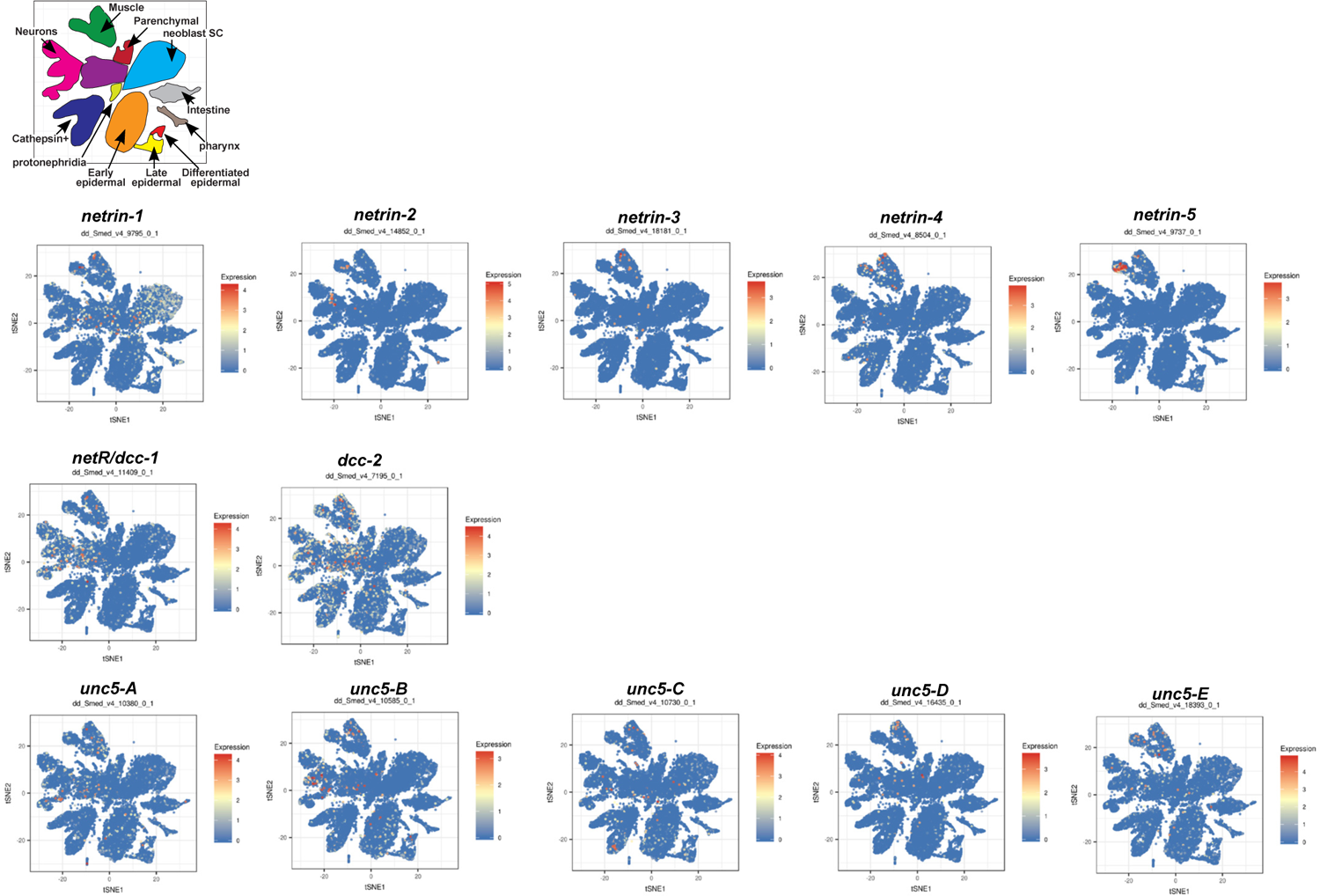
Expression of netrins and netrin receptors from cell atlas. Plots of Netrin and Netrin receptor gene expression from a planarian cell atlas (Fincher et al., 2018; digiworm.wi.mit.edu). Top cartoon shows cluster identities. Netrins and receptors have expression in muscle cells and also in other cell types.

**Figure S3.**
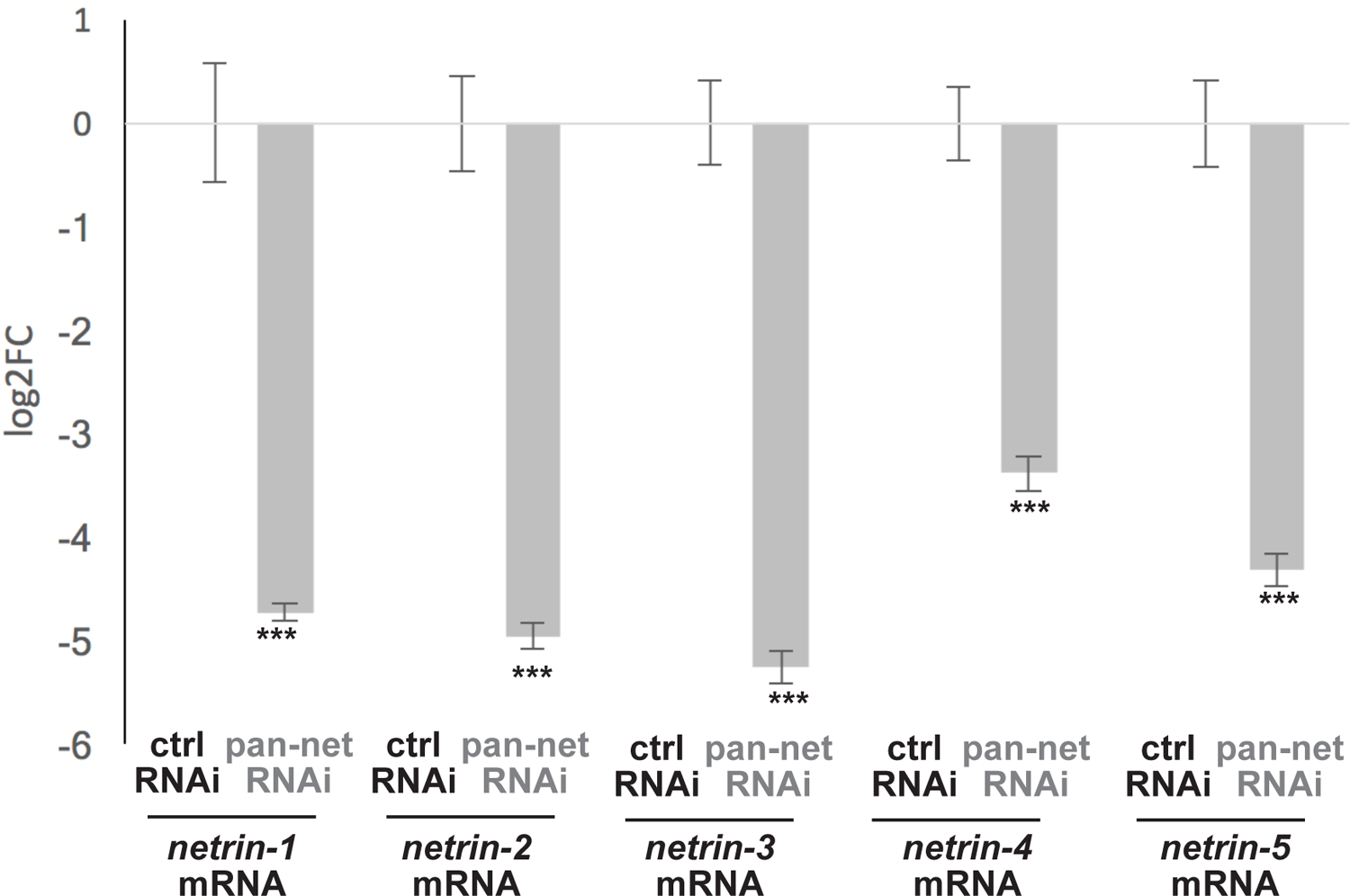
qPCR validating knockdown. dsRNAs targeting netrins −1, −2, −3, −4, and −5 were mixed together and delivered to animals by feeding 6 times over 2 weeks (“pan-net”) and compared to animals treated with control dsRNA. RNA was harvested and qPCR conducted to measure knockdown of each Netrin factor as indicated, using *clathrin* transcript as a normalizing control. Bars show standard error of 4 replicates, ***P<0.001 by 2-tailed unpaired t-test.

**Figure S4.**
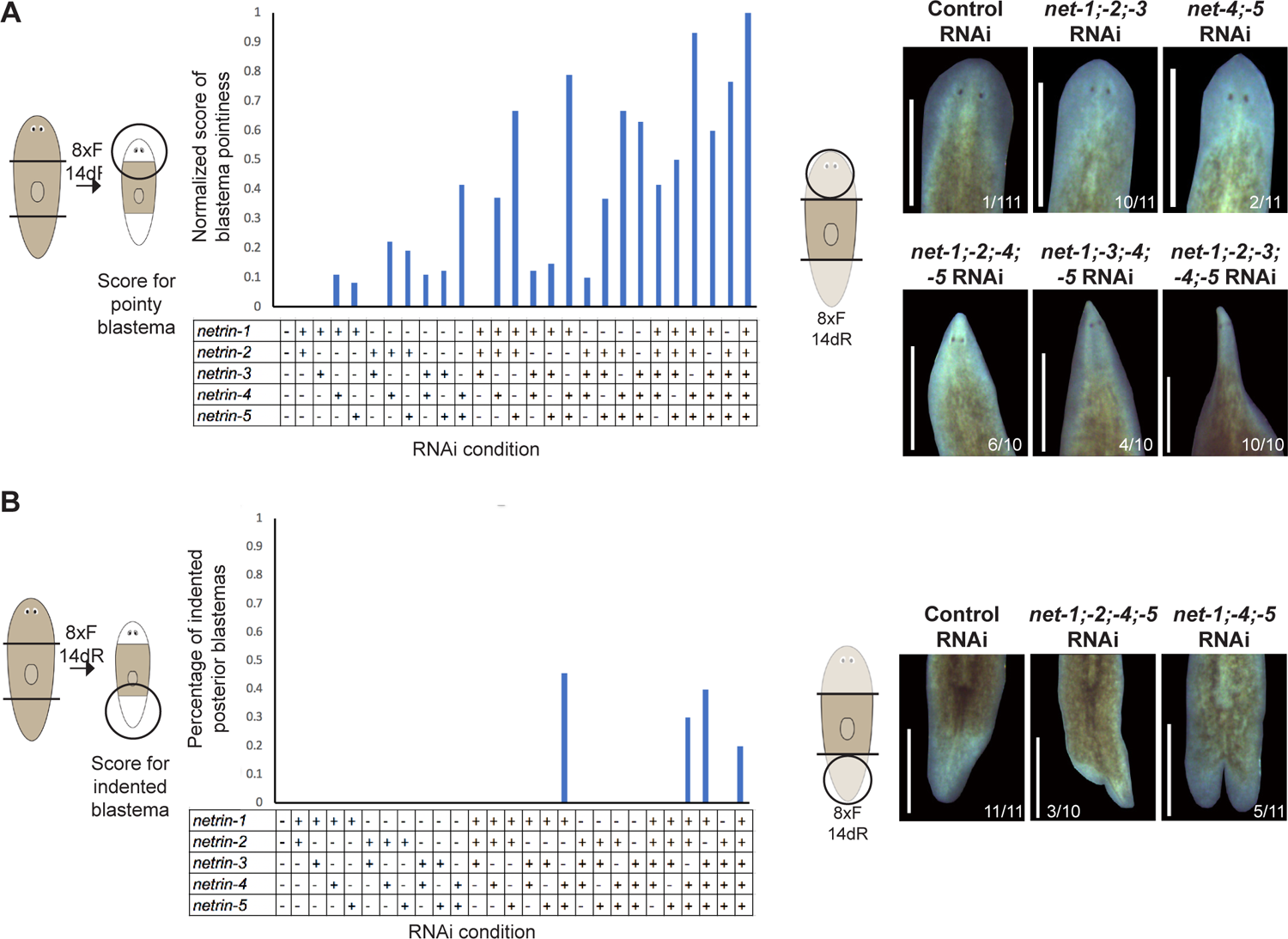
Netrins −1, −2, −4, and −5 promote proper anterior blastema morphology while *net-2, −4,* and *-5* promote posterior blastema morphology. (A) Graph showing regeneration score of anterior blastemas for all combinations of Netrin RNAi. n=7-10 animals. Animals were qualitatively scored based on the degree of pointedness from P0 (normal), P1 (subtle), P2 (pointy), to P3 (highly pointed). In order to compare different conditions with varying penetrance and expressivity, a weighted average phenotype strength score was computed for each treatment by summing the numbers of animals scored in each category multiplied by a scalar of increasing size for higher expressivity (i.e., score = [(# animals in P1) + 2 x (# animals in P2) +3 x (# animals in P3)]/(total number scored)). Average phenotype strength scores were normalized to the value from inhibition of all five Netrins for comparison. Right-Representative images of degrees of blastema pointedness in different combinations of Netrin inhibitions. (B) Graph showing percentage of indented posterior blastemas for all combinations of Netrin RNAi. n=7-10 animals. Right-Representative images of degrees of indented-ness in different combinations of Netrin inhibitions. (C) Live images showing *netrin-1;-2;-3;-4;-5* are not required for blastema formation in regenerating fragments lacking midline.

**Figure S5.**
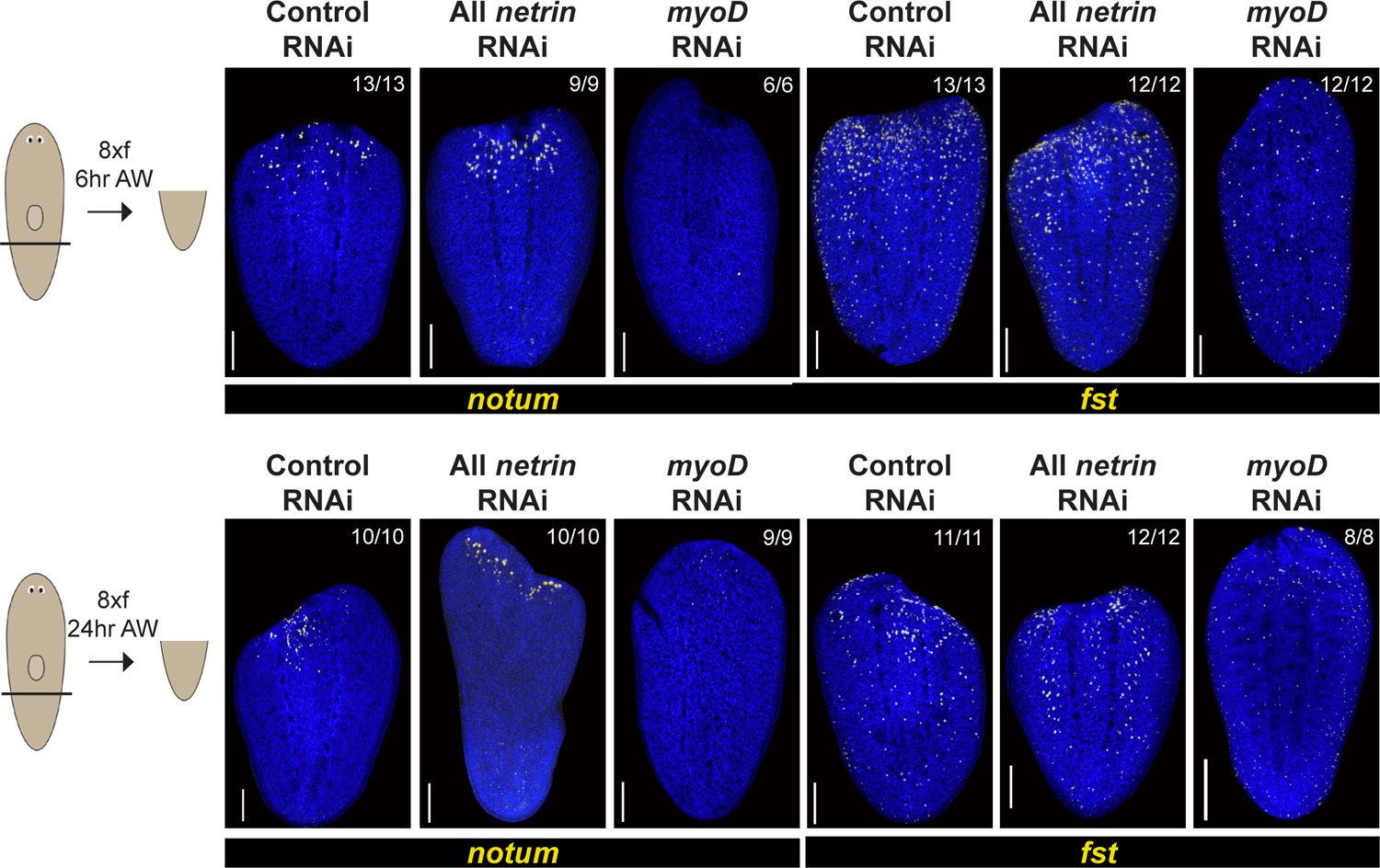
*netrins* are not required for wound-induced expression of *notum* and *follistatin*. **(A)** Animals were fixed following 8 dsRNA feedings and fixed either 6 or 24 hours after injury. FISH showing *notum* or *fst* expression is unchanged in All Netrin RNAi. n=>10 animals. In panel A bars are 200 microns.

## Materials and Methods

### Experimental Model

Asexual *Schmidtea mediterranea* animals (CIW4 strain) were maintained in 1x Montjuic salts between 18–20°C. Animals were fed a puree of beef liver and starved for at least 7 days before experiments.

### In situ hybridizations

Animals were fixed and bleached as described previously (Pearson et al., 2009; King and Newmark, 2013). Riboprobes (digoxigenin- or fluorescein-labeled) were synthesized by in vitro transcription (Pearson et al., 2009; King and Newmark, 2013). Antibodies were used in TNTx/10% horse serum (Roche, Basel Switzerland) at a concentration of 1:2000 for anti-DIG-POD (Roche, Basel Switzerland), and 1:2000 for anti-FL-(Roche), or 1:4000 for anti-dig-alkaline phosphatase used in colorimetric in situ hybridizations (Roche, Basel Switzerland). For multiplex FISH, peroxidase conjugated enzyme activity was quenched between tyramide reactions by sodium azide treatment (100 mM in 1xTNTx) for 45 min at room temperature. Nuclear counterstaining was performed using Hoechst 33342 (Invitrogen, 1:1000 in 1xPBSTx for 45 minutes). Colorimetric in situ hybridizations were developed using NBT/BCIP as described previously (Lander and Petersen, 2016). Primer sequences used for cloning to generate riboprobes are listed in Table S2.

### Immunostaining

Animals were fixed the same as in FISH (King and Newmark, 2013). Bleaching was performed in 80% methanol, 20% hydrogen peroxide on a lightbox overnight. Following rehydration washes, animals were washed and rocked in PBSTx/10% horse serum (Roche, Basel Switzerland) 8 times for 6 hours. Animals were incubated in the anti-muscle mouse monoclonal antibody 6G10-2C7 (Iowa Hybridoma Bank) at 1:1000 overnight, then washed in PBSTx/10% horse serum 8 times for 6 hours. Animals were incubated overnight in an anti-mouse Dig conjugated antibody in PBSTx/10% horse serum. After 8 PBSTx washes, tyramide amplification was performed using commercial tyramide kit. Finally, animals were washed in PBSTx 8 times, 20 minutes each.

### RNAi administration

RNAi by feeding was performed in vitro transcribed dsRNA mixed with liver paste and 5% red food dye (Rouhana et al., 2013). For single RNAi experiments, dsRNA at a concentration of 2-4 ug/ul was mixed with liver in a 1:5 ratio. In RNAi experiments with multiple dsRNAs, each dsRNA was mixed with liver for a total concentration of 1:10 for each dsRNA. In all RNAi experiments animals were fed RNAi food every 2-4 days for the length of experiment indicated. For homeostatic experiments, animals were starved for at least 5 days before fixing. In regeneration experiments, animals were amputated the same day of the last feeding, or in Figure 1C, 1D, Figure S4 at 5 days after the final RNAi feeding. dsRNA targeting the *Photinus pyralis* luciferase (not present in the planarian genome) was used as a negative control RNAi condition. Primers sequences used for RNAi are listed in Table S2.

### Image Acquisition and Quantification

Live images of animals were performed with a Leica M210F dissecting scope with a Leica DFC295 camera. FISH images were performed a Leica DM5500B compound microscope with optical sectioning by Optigrid structured illumination or a Leica Stellaris or Leica TCS SPE confocal compound microscopes. Fluorescent images collected by compound microscopy are maximum projections of a z-stack and adjusted for brightness and contrast using Metamorph, Adobe Photoshop, ImageJ, or Leica LAS.

### qPCR

RNA was harvested from 4 biological replicates of 4 animals per condition using a Turrex tissue homogenizer. cDNAs were synthesized using oligo-dT and Superscript III reverse transcriptase (Life Technologies) for 1 hour at 50°C. EvaGreen qPCR mastermix (Midwest Scientific BEQPCR-S) was used for amplification and detection. Per-sample log2-fold change expression were determined by the delta-delta Ct method, normalizing to expression of housekeeping gene clathrin probe set control and to RNAi negative controls targeting *Photinus pyralis* luciferase (not present in planarian genome). Statistical significance was determined through unpaired two-tailed t-tests comparing ddCt values between control RNAi and target RNAi conditions. Primers for qPCR: *netrin-1* (5’-taaagagcatcagggtctatgtga-3’, 5’-ataatatcgacgcttcaatttcca-3’); *netrin-2* (5’-gaattggtatgacaagacaactgc-3’, 5’-gttttgcgtcgcatatactacaat-3’); *netrin-3* (5’-cgaaaccctatgtaaaaagaagga-3’, 5’-cccttggagattttataggacaga-3’); *netrin-4* (5’-cctgttaaattgaaatcgctctct-3’, 5’-aataaaataatttggggatgctga-3’); *netrin-5* (5’-tgcacaaaatgctacatttgaagt-3’, 5’-atcctttctgaccgcatgtact-3’); *clathrin* (5’-agccgcatgtagttggttagtg-3’, 5’-agtctacagaaaaccatcaattgtttcaaa-3’).

### scRNAseq differential gene expression testing

We mined a prior single-cell RNAseq atlas generated by DropSeq (Fincher et al., 2018) and recovered 15 *dd23400*+*slit*+ cells (at least 1 read for *bambi-2/dd23400* and at least 2 reads for *slit*) and compared these to 143 randomly selected muscle cells (at least 7 reads for muscle marker *collagen* dd840 and no reads for *dd23400* or *slit*) and analyzed for differential gene expression using the Single Cell Differential Gene Expression (scde) method (Kharchenko et al., 2014). Table S1 shows results of differential testing along with identity of cells selected for analysis.

### Phylogenetic analysis

Bayesian inference phylogenetic tree was generated from alignment with MUSCLE and calculated with MrBayes v3.2.3 using a GTR substitution model with gamma-distributed rate variation across sites and a proportion of invariable sites. Runs of 50000 generations converged with an average standard deviation of split frequencies <0.01 and 25% of trees were discarded as burn-in.

## Notes

### Competing Interest Statement

The authors have declared no competing interest.

### Summary of Updates

Text and abstract revised; Figure 2 removed;

